# Did Energy Costs of Walking Limit the Evolution of a Larger Human Birth Canal?

**DOI:** 10.1101/2024.12.11.627886

**Authors:** Ekaterina Stansfield, Clara Egner, Philipp Mitteroecker, Hans Kainz

**Affiliations:** Department of Evolutionary Biology, University of Vienna, Austria; Department of Sport and Human Movement Science, University of Vienna, Austria

**Keywords:** forward motion modelling, cost of transport, human evolution, obstetrical dilemma, leg length, pelvis form

## Abstract

Energetic disadvantages of a wider pelvis have long been central to explanations of the small human birth canal and the relatively risky birth process, iconic for antagonistic selection in evolution. While static mechanics predict such effects, recent empirical studies question this view. By altering a generic musculoskeletal model beyond modern human variation one parameter at a time and applying predictive forward modelling of human gait, we examined how body dimensions impact locomotion efficiency and inferred selective pressures on past morphologies. Our findings confirm that a wider pelvis increases abductor muscle demands, as lever mechanics suggest. However, dynamic walking compensates via other muscles, minimizing the total transport cost. Moreover, slightly longer legs easily offset the modest energetic costs of a larger birth canal. We propose that other factors, such as pelvic floor stability, likely constrained the evolution of the human birth canal, rather than energetic disadvantages alone.

## Introduction

The relatively small human birth canal and the associated risk of obstructed labour have prompted numerous hypotheses about selective forces that counteract the obvious advantages of a larger birth canal for parturition ^1–4^. Also, the pronounced sex differences in pelvic form^5–7^ may imply a selective force in men, such as locomotor efficiency or male reproductive function, that counteract birth-related selection in women ^8–10^. For many decades, biologists thus viewed pelvic morphology as an evolutionary compromise between the partly antagonistic demands of childbirth and bipedal locomotion ^1–3,11^. The idea of such a functional and evolutionary conflict was supported by inferences from static lever mechanics, allowing Lovejoy et al. ^12^, Lovejoy ^13^, and Tague and Lovejoy ^14^ to observe that wider hips increase the lever arm of the body’s centre of mass to the hip joint centre and therefore increase abductor muscles’ energy expenses. Broader pelves would thus be energetically more costly. These static analyses, however, neglected the dynamics of the walking pattern, which constantly changes the centre of mass position and lever arms, length and contraction velocity of muscles, which all affect muscle forces^15^ and, therefore, energy expenses^16^.

More recent empirical studies measured metabolic energy to evaluate the effects of a broader pelvis on bipedal locomotion. Contrary to the classic expectations, Warrener et al. ^17^ found no link between the mediolateral dimensions of the pelvis and metabolic energy consumption in young recreational runners. Gruss et al. ^18^ experimentally demonstrated that subjects with wider pelves take longer strides for a given velocity. They flex and extend their hips less, suggesting a smoother pathway of the centre of mass (COM). In another kinematic study, Witcombe et al. ^19^ showed that females exploited the breadth of their obstetric pelvis to obtain longer strides relative to leg length. Further studies of locomotion in females asserted their efficiency when carrying loads and crossing rugged terrains ^20^.

Experimental studies are affected by many confounding factors, such as body weight ^21^, walking pattern ^22^, walking speed ^23^, muscle coordination ^24^, muscle properties ^25^, and musculoskeletal geometry ^26,27^, which can all differ between individuals and affect muscle forces and energy consumption. Moreover, much of the discussion about the impact of pelvis form on locomotion efficiency has focused on whether women are more or less efficient than men and whether they are ‘disadvantaged’ due to the evolutionary pressure from obstetrics ^28,29^. However, it has proven challenging to pinpoint any differences in energy expenses due to sex differences in pelvis shape. For example, on average, women are shorter than men and require less energy to move and maintain their basal metabolism. Still, no sex differences were found in daily energy expenditure if age, fat-free and fat mass were considered ^30^. Women were reported to show greater energy economy in running ^31^ and walking ^32,33^. Still, sports science studies do demonstrate that the geometry of the musculoskeletal system affects muscle forces and sports performance ^34,35^. Overall, it is very difficult to control for all these covariates experimentally or statistically when relating pelvis shape to energy consumption, and it is impossible to disentangle these many factors when comparing only two sex means.

Here, we used an in-silico approach to investigate how different pelvic dimensions affect locomotion efficiency and energetics. We used a predictive forward modelling framework of the human gate in a generic musculoskeletal model (the “base model”) with 92 musculotendon actuators of the lower extremities and torso^36^. The body dimensions of the base model were changed one parameter at a time by +/- 50%, which is roughly a threefold extrapolation of modern human variation. This way, we could gauge the effect that a single anatomical dimension might have on locomotion energetics, independent of biometric parameters that would covary in experimental settings. Extrapolating modern human variation allowed us to infer past selective pressures on morphologies no longer present in recent populations.

We hypothesised that a broader pelvis results in a higher cost of transport (COT) when all other variables are kept the same. We compared the effect of pelvis breadth with an equivalent impact of pelvis depth (i.e., the anteroposterior diameter, both with and without a change in torso dimensions) and leg length on the total COT and the COT for the abductor muscles (Fig. 1). All other body dimensions, mass, and mass distribution were kept constant. The speed was set at 1.33 m/sec for all models. Predictive simulations were run to minimise a cost function that included metabolic energy rate, muscle activity, joint accelerations, passive joint torques, and arm excitations as parameters. We used kinematics obtained from a motion capture trial as the initial guess (‘hot start’) ^36^.

**Figure 1.**
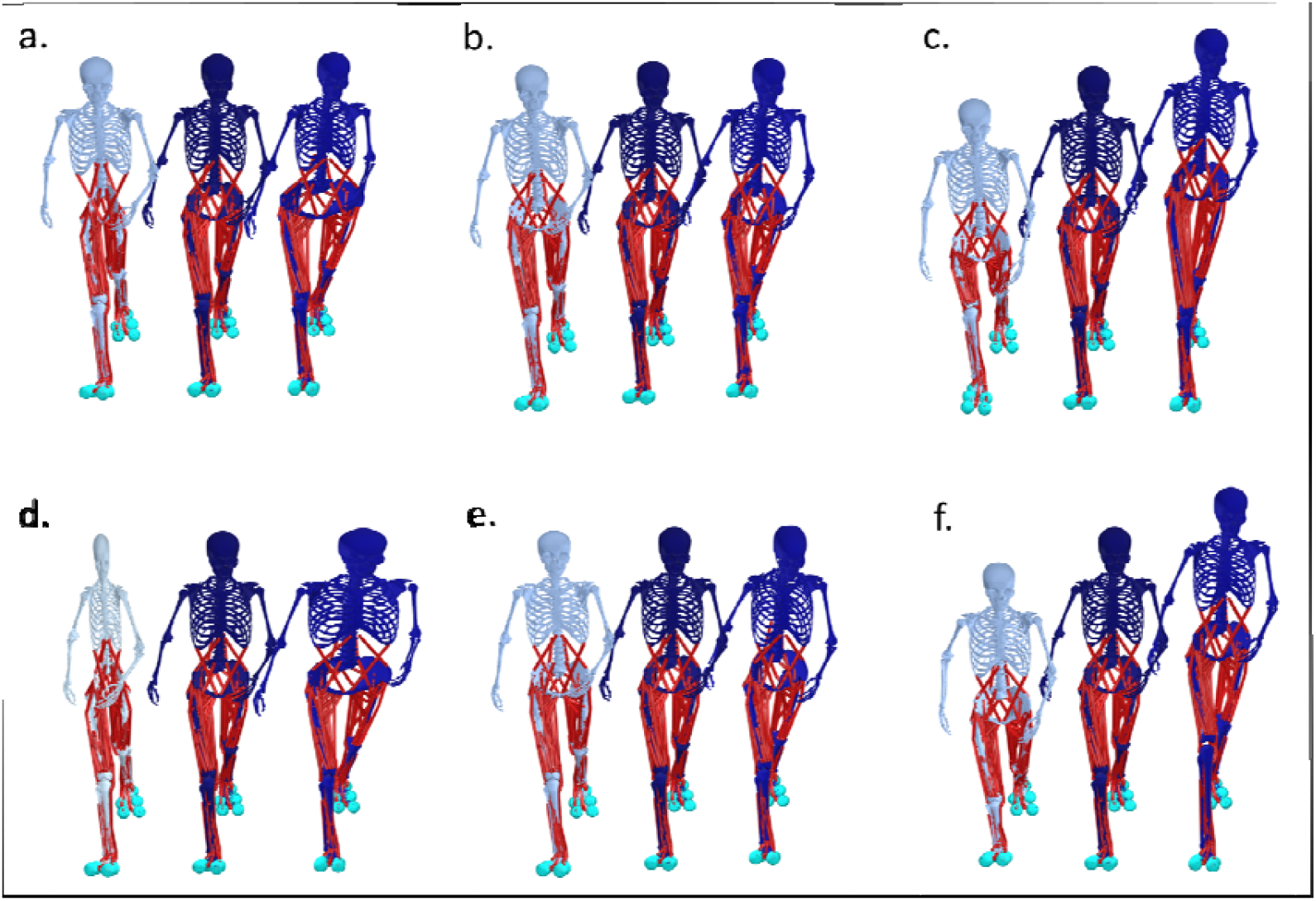
Examples of the scaled model, with the lower and upper extremes shown together with the base model: (a) pelvis width of 0.5, base model, and pelvis width 1.5; (b) pelvis depth 0.5, base model, and pelvis depth 1.5; (c) femur length 0.6, base model, and femur length 1.4; (d) pelvis and torso width 0.5, base model, pelvis and torso width 1.5; (e) pelvis and torso depth 0.5, base model, pelvis and torso depth 1.4; (e) tibia length 0.6, base model, and tibia length 1.4.

## Results

All models whose kinematics were successfully computed produced human-like walking gait with somewhat similar kinematics (Fig. S1). Models with very wide hips tended to have elevated pelvis tilt. Models with deeper pelvis also tended to have a higher magnitude of lumbar extension. Models with the shortest legs had more variation in the ankle angle over the predicted stride. However, they still produced the expected human heel strike.

### Total Cost of Transport (COT)

The total COT was affected by some, though not all, body dimensions (Fig. 2). The increase in pelvis width produced only a small increase in the COT. When pelvic width was modified jointly with torso width, basically no effect on the COT was visible. Pelvis depth had a larger impact on the COT than pelvis width, regardless of whether the changes occurred only in the pelvis or the pelvis and torso. Effects on the COT were strongest for femur and tibia lengths. An increase in femur or tibia length (while keeping the other length constant) reduced the COT, but an increase in leg length beyond +20% did not produce any further energy benefits. Solutions for models with femur lengths -50% and +50% did not converge. When expressing the range of COT values calculated for one dimension relative to the base COT (Table 1), femur and tibia lengths showed an effect of approximately 18% of the base COT, pelvis depth less than 5%, and pelvis width 2.8%.

**Table 1.** Range of total COT values in the tested models calculated by subtracting the minimum COT observed in an experiment from the maximum and expressing the difference as a percentage of the base model’s COT. Due to the discrepancy in the number of models per experiment, we only report the range of COT values for the parameter range from 0.6 to 1.4 of the base model.

|  | pelvis width | pelvis torso width | pelvis depth | pelvis torso depth | femur length | tibia length |
| --- | --- | --- | --- | --- | --- | --- |
| Range<br>(J/m/kg) | 0.093 | 0.043 | 0.154 | 0.162 | 0.76 | 0.731 |
| Range as per cent of base | 2.2 | 1 | 3.7 | 3.9 | 18.4 | 17.7 |

**Table 2.** Ranges for the COT of the abductor muscles (from 0.6 to 1.4 factor). The values were calculated by subtracting the minimum observed COT from the maximum in each experiment and expressing the difference as a percentage of the base model’s COT.

| name | pelvis_width | pelvis_torso_width | pelvis_depth | pelvis_torso_depth | femur_length | tibia_length |
| --- | --- | --- | --- | --- | --- | --- |
| Range (J/m/kg) | 0.128 | 0.128 | 0.038 | 0.042 | 0.07 | 0.096 |
| Range as per cent of base | 26.%1 | 26.1% | 8.1% | 8.1% | 14.3% | 19.6% |

**Figure 2.**
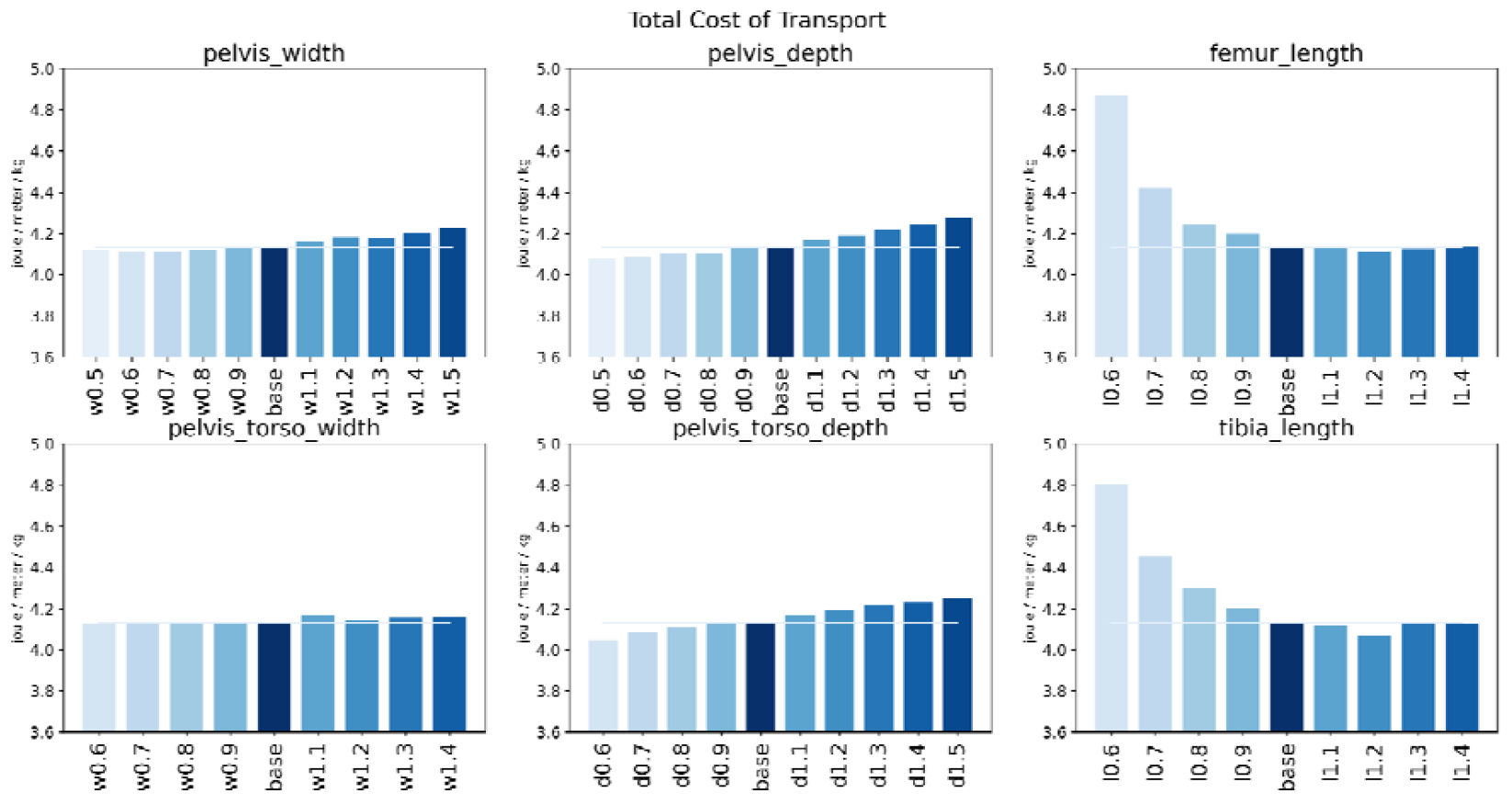
The total cost of transport (COT) per model for one walking cycle and a range of different body dimensions. Each panel depicts a simulation where we varied one of the geometric parameters in steps of 0.1 multiples of the base model. Lighter bars represent smaller values of the parameter. The darkest colour indicates the base model.

### Variation in the cost of transport (COT) per muscle

Focusing on individual muscles, one can observe differential effects on their COT resulting from changes in gross morphology (Fig. 3). In models with increased pelvis width, the total COT rose for gluteus maximus 1 and 2, gluteus medius 1, 2, and 3, and gluteus minimus 1 and 2, with a particularly high increase for the internal oblique muscle. The COT decreased for gluteus maximus 3, adductor magnus 1, 2, and 3, iliacus, and psoas.

**Figure 3.**
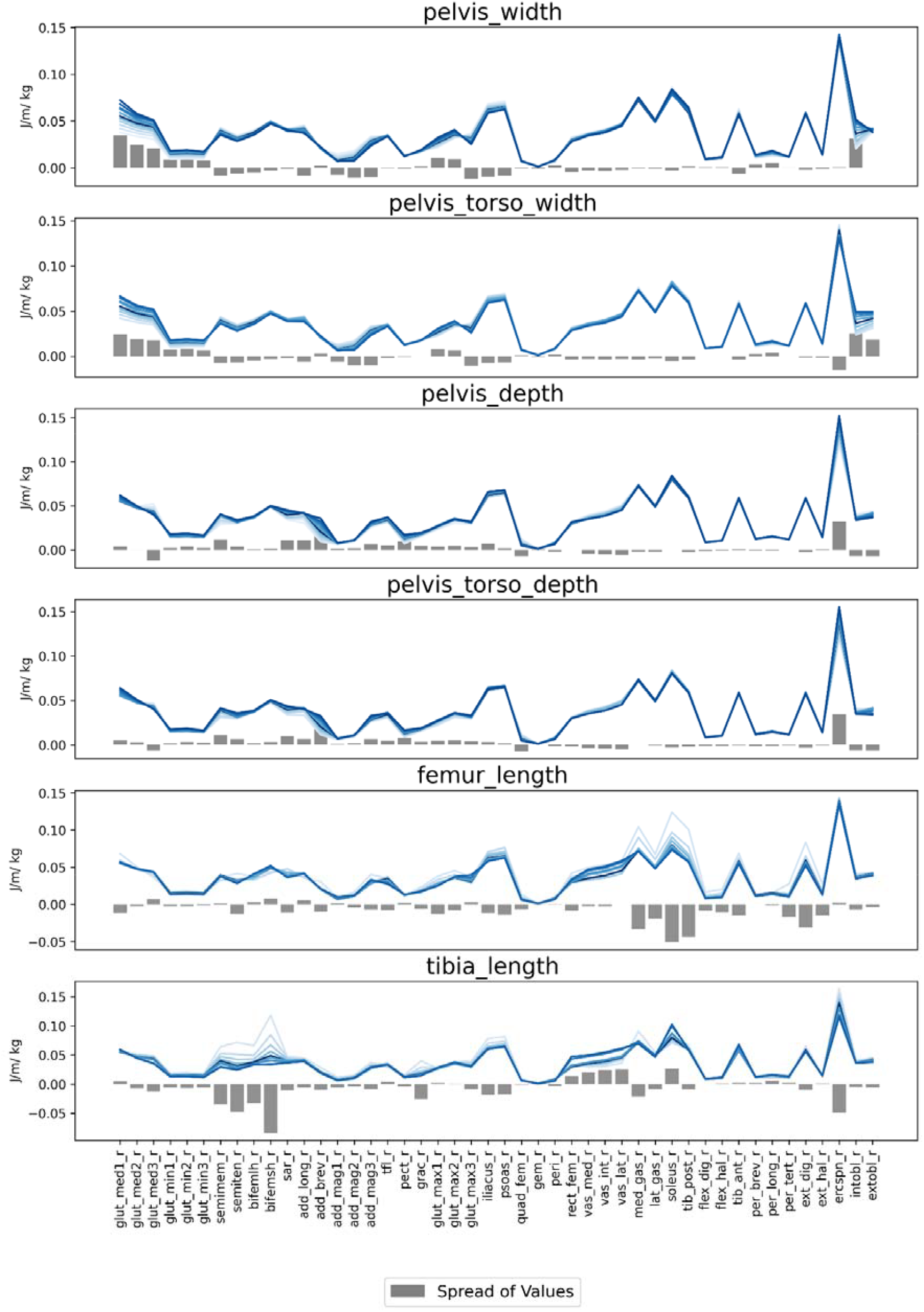
COT per muscle. As in Fig. 2, each plot shows an experiment that changed one of the geometric parameters. Lighter colours express smaller values of the parameter. The darkest colour indicates the base model. Grey bars show the magnitude and direction of change across models as the value of the varied parameter (width or length) increases from the minimum to the maximum.

In models where both pelvis and torso width were modified, the pattern was similar to that of pelvis-only changes, but the COT for the erector spinae decreased while the COT for the external oblique increased. In other words, as the mediolateral dimensions of the pelvis and torso increased, the COT of laterally positioned muscles increased, while that of medially positioned muscles decreased.

When pelvis depth increased, the COT rose for the semimembranosus, sartorius, adductor longus, adductor brevis, pectineus, iliacus, and erector spinae muscles but decreased for gluteus medius 3, quadratus femoris, and both the internal and external oblique muscles. When torso depth increased along with pelvis depth, the overall COT pattern remained largely unchanged, except for the gluteus medius 3, which no longer exhibited a significant increase in COT.

As leg length increased, total COT decreased from the smallest leg length to the base length by +20% (Fig. 1). However, the load distribution varied depending on whether the proximal or distal segment of the leg was altered. Increased femur length reduced the COT for the medial and lateral gastrocnemius, soleus, tibialis posterior, peroneus tertius, extensor digitorum, and extensor hallucis muscles. However, longer femora also led to higher COT in some models for the vastus muscles, though the pattern was inconsistent. A longer tibia significantly reduced the COT for the semimembranosus, semitendinosus, biceps femoris (both long and short heads), gracilis, iliacus, psoas, and medial gastrocnemius muscles. The erector spinae also benefited from a longer shank. COT increased for the rectus femoris, vastus, and soleus muscles.

### COT for abductor muscles

Abductor COT increased almost linearly with a broader pelvis (Fig. 4). An increase in pelvis depth within the +/- 30% range from the base also led to a rise in the COT of the abductor muscles. Increasing leg length resulted in a decrease in abductor muscle COT. However, this effect was observed only within a range of -10% to +40% from the base for femur length and -20% to +40% for tibia length. The range of COT values was largest for pelvis width, with changes reaching up to one-third of the base abductor COT, followed by tibia and fibula lengths and pelvic depth (Tab. 2).

**Figure 4.**
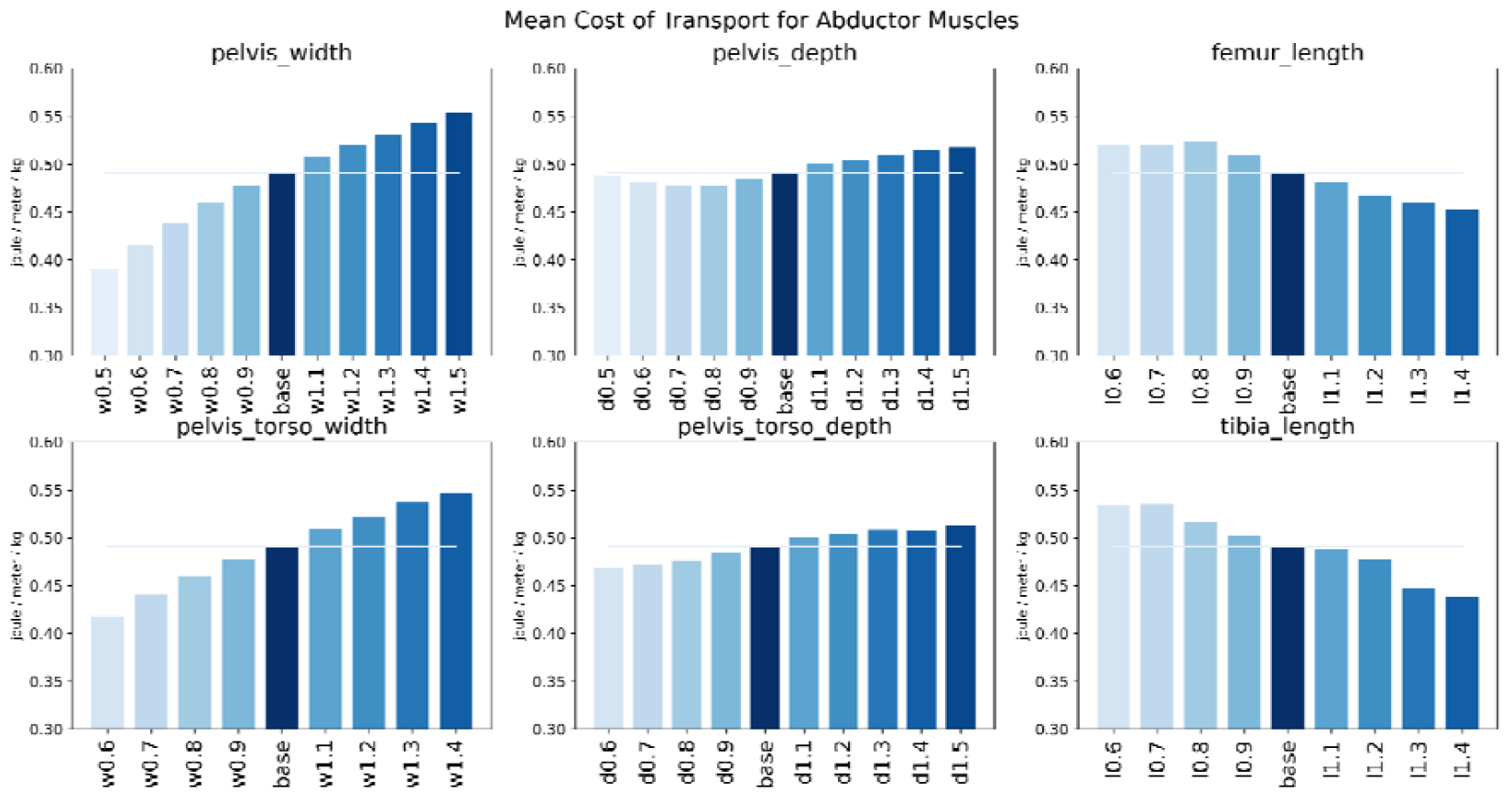
Total COT for the abductor muscles per one walking cycle. Each plot depicts an experiment where we vary one of the geometric parameters of the model.

## Discussion

The putatively negative effect of pelvic width on locomotion efficiency has long been considered the main evolutionary cause of the narrow human birth canal and has been iconic for antagonistic selection in human evolution. Experimental and in silico tests of this assertion have only been conducted in the last decade and have led to largely contradictory results. Here, we tested this assertion with a detailed predictive forward modelling framework, which allowed us to assess the effect of pelvic breath independent of other body dimensions.

In contrast to Kramer and Silvester ^37^ and Warrener et al. ^17^, who analysed motion capture data, our simulations demonstrated a substantial impact of pelvis width on the energy demands of the abductor muscles. This discrepancy is due likely to the difficulty in controlling confounding factors in experimental motion capture and oxygen consumption studies. However, when considering the total cost of transport (COT) for all simulated muscles, pelvis (and torso) width had only a small effect. Due to the redundancy of the human musculoskeletal system ^38^ and the dynamics of walking, other muscle groups compensated for the increase in the abductors’ COT, resulting in little effect on total COT. The limited impact of pelvis width on total COT aligns with empirical findings by Warrener et al. ^17^. We did not observe longer strides in models with wider hips in the scaled models as did Gruss et al. ^18^ and Whitcome et al. ^19^ in motion capture studies (Fig. S2).

Pelvic depth affected various muscles and even had a greater impact on total COT than pelvic width. This is consistent with earlier suggestions on how the anteroposterior diameter of the human pelvis affects locomotion, particularly in relation to the distance between the sacroiliac and hip joints and its effect on pelvic tilt and lumbar lordosis ^1,2,9,39–41^.

In our simulations, increasing femur or tibia length affected different muscle groups. It led to a major decrease in total COT, which brings forward the importance of leg length in walking. The rate of COT reduction was similar for both leg segments till up to 20% of their base size. The stable COT for more than a 20% increase in leg length likely owes to the constant speed of 1.33 m/s in all models. For longer legs, this may not be the optimal speed, thus requiring more energy ^42^.

Negative correlations between leg length and energy consumption during walking and running are well documented. For example, Van der Walt and Wyndham ^43^ proposed a regression equation where oxygen consumption in locomotion is positively related to mass and the square of velocity but negatively related to leg length. Silva and Machado ^44^ calculated that for a simple bipedal robot moving in the sagittal plane at a constant speed, minimal power requirements occurred when the leg length was 85% of total height, and the ratio between proximal and distal limb segments was 0.35:0.65. Similarly, Steudel-Numbers and Tilkens ^45^ demonstrated a negative correlation between limb length and oxygen consumption in humans walking at different speeds on a treadmill. Bereket ^46^ further suggested that shank length significantly influenced oxygen consumption once speed and mass effects were controlled.

This study is limited by the ability of our model to reflect real-world conditions. Falisse et al. ^36^ performed a sensitivity analysis on the original model, adjusting various parameters to produce realistic gait patterns in healthy and impaired individuals. Simulated walking patterns aligned well with empirical data. We used the recommended cost function parameters and settings for a healthy, bilaterally symmetric subject, ensuring our model accurately reproduces human walking gait patterns.

In our experiments, we intentionally altered only one parameter at a time, which does not reflect realistic covariation patterns in the musculoskeletal system. Moreover, the range of alterations was much wider than the typical range of human variation, and these model results thus are extrapolations that cannot easily be validated. Lastly, this study did not address other biomechanical effects of pelvis geometry, such as joint loading and injury risk. Further research is needed to confirm our findings and explore biomechanical limitations imposed on the human musculoskeletal system by bipedal locomotion.

In conclusion, the energetic efficacy of bipedal locomotion has long been the prime explanation for the selection towards a small birth canal, but a series of recent studies have challenged this reasoning ^17,18,29,37^. Our findings confirm that a mediolaterally wider pelvis indeed increases the energetic demands of abductor muscles, as previously predicted by static lever mechanics^12,13,14^. However, in the dynamic walking process, other muscles compensate for these demands so that the total costs of transport only weakly increase with pelvic breath. This weak association was shown in the reanalyses of data of Warrener ^17^ by Haeusler et al. ^9^, and of Kramer and Silvester ^37^ by Grunstra et al. ^10^. The cliff-edge model of obstetric selection^47,48^ predicts that only weak selection for a small birth canal suffices to explain the observed rates of obstructed labour. Yet, it seems unlikely that the minor effects of pelvic width on total COT are indeed the main evolutionary drivers behind the small human birth canal. More strongly, the total COT increases with pelvic depth. Moreover, the results of the present study suggest that slightly longer legs could easily compensate for the relatively weak energetic costs of a larger birth canal.

Even though total COT seems to impose only weak selective pressure on pelvic width, other aspects of bipedal locomotion, which we did not assess here, may play a role. For example, the excessive engagement of the abductor muscles of the hip may affect an individual’s endurance capacity and, indirectly, their reproductive fitness in an environment where endurance is required for survival. Moreover, a larger pelvis would be linked with larger trunk diameters and larger body mass, which may require more energy for maintaining basic metabolic functions. It is also well documented that body proportions and, thus, pelvic width are important for heat dissipation and have adapted to the local thermal environment ^49–51^.

Overall, our results indirectly support the pelvic floor hypothesis as an alternative evolutionary explanation for the tight foeto-pelvic fit in modern humans. This hypothesis suggests that a larger birth canal would reduce the stability of the pelvic floor and thus increase the risk of disorders such as incontinence and pelvic organ prolapse ^8,52,53^.

Epidemiological, morphometric, and biomechanical studies ^41,54–57^ support the pelvic floor hypothesis much more clearly than the classic “obstetrical dilemma” hypothesis of locomotion efficiency.

## Material and Methods

### General Set-up

We used a predictive modelling framework developed by Falisse et al. ^36^ in Python to model the gait of a healthy person using 61 models, including:

1. A base model driven by 92 musculotendon actuators of the lower extremities and torso, with five degrees of freedom at each lower extremity and three degrees of freedom at each hip ^58^,
2. The same model as the base altered but one of the following six dimensions was altered incrementally from the factor 0.5 to the factor 1.5:
  - pelvis width,
  - pelvis depth,
  - pelvis and torso width,
  - pelvis and torso depth,
  - femur length, and
  - tibia length.

This range of alterations widely exceeds the typical range of human variation. For instance, bi-iliac breadth in American women ranges from 23.9 mm at the 5th percentile to 34.9 mm at the 95th percentile and from 25.3 to 34.5 mm in men^59^. This range is about +/-14% of the women’s average of 29.1 mm. For men, the range corresponds to +/-13% of the average of 31.9 mm. The upper leg length is an average of 42.3 mm in men, with 90% of the population fitting in the range between 37mm and 47.8 mm, which corresponds with +/-13% of the mean. In women, the range is from 33 mm to 43.9 mm, corresponding to +/-14%of the mean (38.6 mm). All other geometry, mass and mass distribution were kept constant. The speed was set at 1.33 m/sec for all models.

Predictive simulations were run to minimise a cost function that included metabolic energy rate, muscle activity, joint accelerations, passive joint torques, and arm excitations as parameters. The optimal control problem was solved with the help of the direct collocation method^60,61^, with 50 frames calculated over the one-stride period for each model. The metabolic energy rate for each muscle was obtained with the help of a smooth approximation of the Bhargava et al.^62^ model of muscle contraction, adapted by Falisse et al.^36^ for the forward modelling. Muscle-specific COT was then the metabolic energy consumed per unit distance travelled. The abductors’ COT was represented by the sum of COT for abductor muscles on one side, while the total COT integrated the metabolic energy of all muscles.

The number of solved models varied slightly across the experiments. A maximum of 11S models were successfully computed in the experiments that altered pelvis width and depth. However, only nine models ran successfully in experiments that adjusted pelvis and torso widths and femur and tibia lengths. In these cases, models with scale factors of 0.5 and 1.5 failed to converge. Ten models worked for pelvis and torso depths, with only one scaled to 0.5 depth factor failing. We compared the effect of the changing geometry on the total COT, COT per muscle and abductor muscles COT by calculating the range of the energy measures given the varied parameters.

### Local solutions

Gait trajectory optimisation is an example of a non-linear problem where finding a global solution is frequently impossible and is replaced by finding a local solution. In other words, a solution is found that minimises the objective function among feasible points that are near it but does not guarantee to have a lower objective value than all other feasible points ^63^. In practice, multiple gait trajectories can be predicted depending on the value of the initial guess or the weights of the optimised parameters, and the quality of the solution can be judged by its proximity to the human gait pattern and by comparison of the value of the optimised parameters between the solutions. Optimal control problems were solved using Python with the help of automatic differentiation using a nonlinear optimiser (CasADi^64^).

We conducted several tests to assess the solutions for the 61 models in the present study, where we used recommended weights for optimised parameters and pre-defined kinematics as an initial guess (‘hot start’^36^). First, we perturbed the values of the weights by +/- 10% and observed the effect on the total COT of 12 out of 61 models (Fig. S3). We then solved the gait prediction problem for each of the 61 models using zeros as the initial guess (‘cold start’). These results are presented in the Figure S4 of the Supplementary Material.

Our solutions were stable across the 10% perturbations of the weights of the optimised parameters. When the cold start was used as the initial guess, however, it produced a variation of predicted gaits. Several models walked on toes, which resulted in a significantly higher total COT compared with the models that were used during the regular walking pattern. These solutions represent alternative local minima, which are less optimal and less realistic than the ones obtained with the ‘hot start’. Therefore, we are reasonably confident that the ‘hot start’ solutions with the recommended weights of the optimised parameters are the best solutions for the system and are closest to the natural human gait.

## Data Availability

Computation results will be made publicly available on publication at https://osf.io/.

## Code Availability

Code has been made available by Antoine Falisse at https://github.com/antoinefalisse/opensimAD and https://github.com/antoinefalisse/predsim_tutorial.

## Acknowledgements

We are very grateful to Dr Antoine Falisse for providing the code for the forward simulation of walking in the open access and for his support in answering our questions during the project development. We are also grateful to Friedl de Groote, who suggested the series of tests we ran to explore the stability of the presented solution. This work was funded by Austrian Science Fund Grant P 35714-B (ES).

## Author contributions

Conception and design: E.S., H.K, P.M.

Analysis of the data: E.S., C.E.

Interpretation of the data: E.S., P.M.

Drafting and revision of the manuscript: E.S, P.M., H.K.

