## Supplementary Material for "Did Energy Costs of Walking Limit the Evolution of a Larger Human Birth Canal?"

**Figure S1.** Kinematics of models, sorted by the parameter that was varied in each experiment.

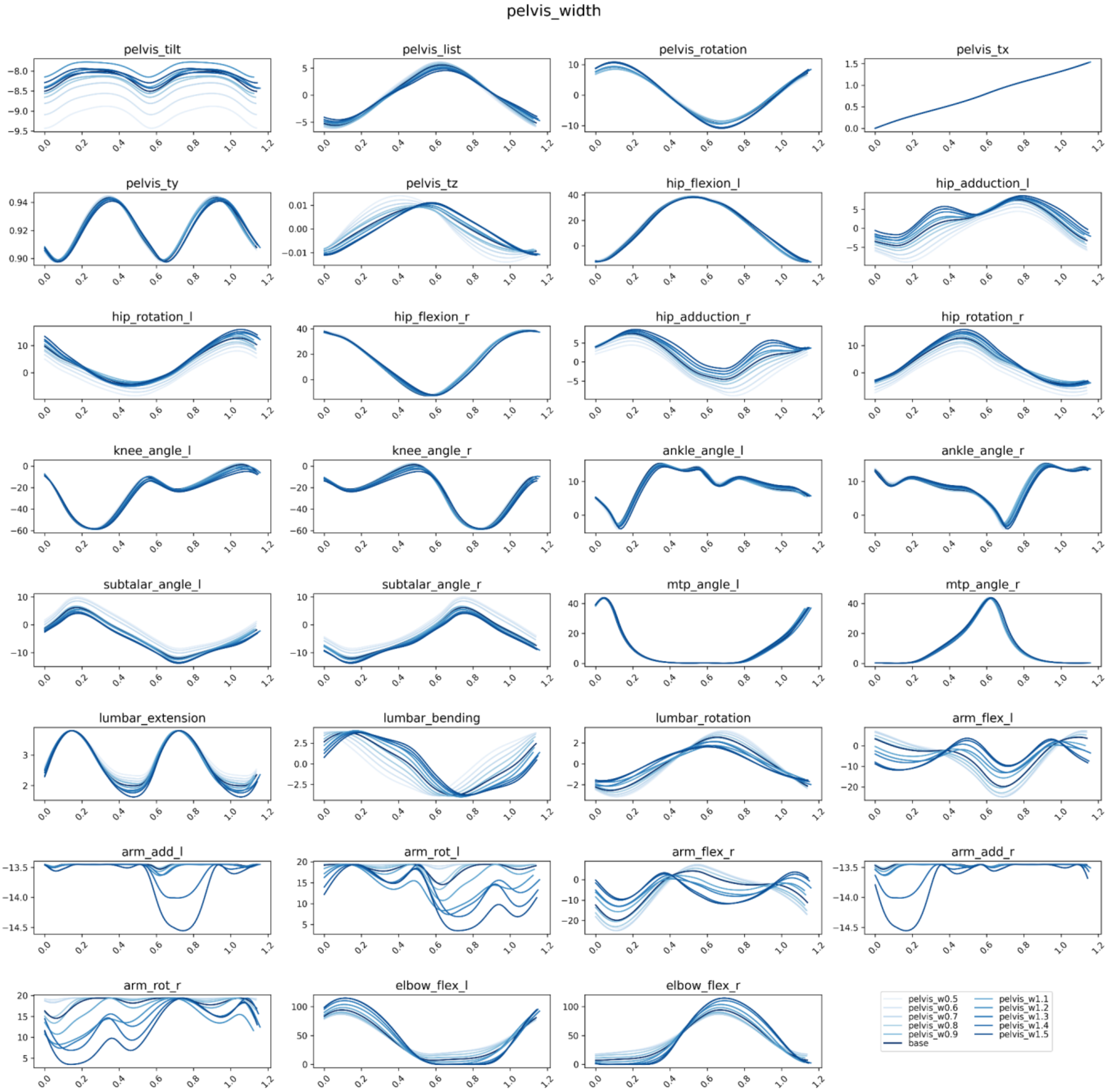

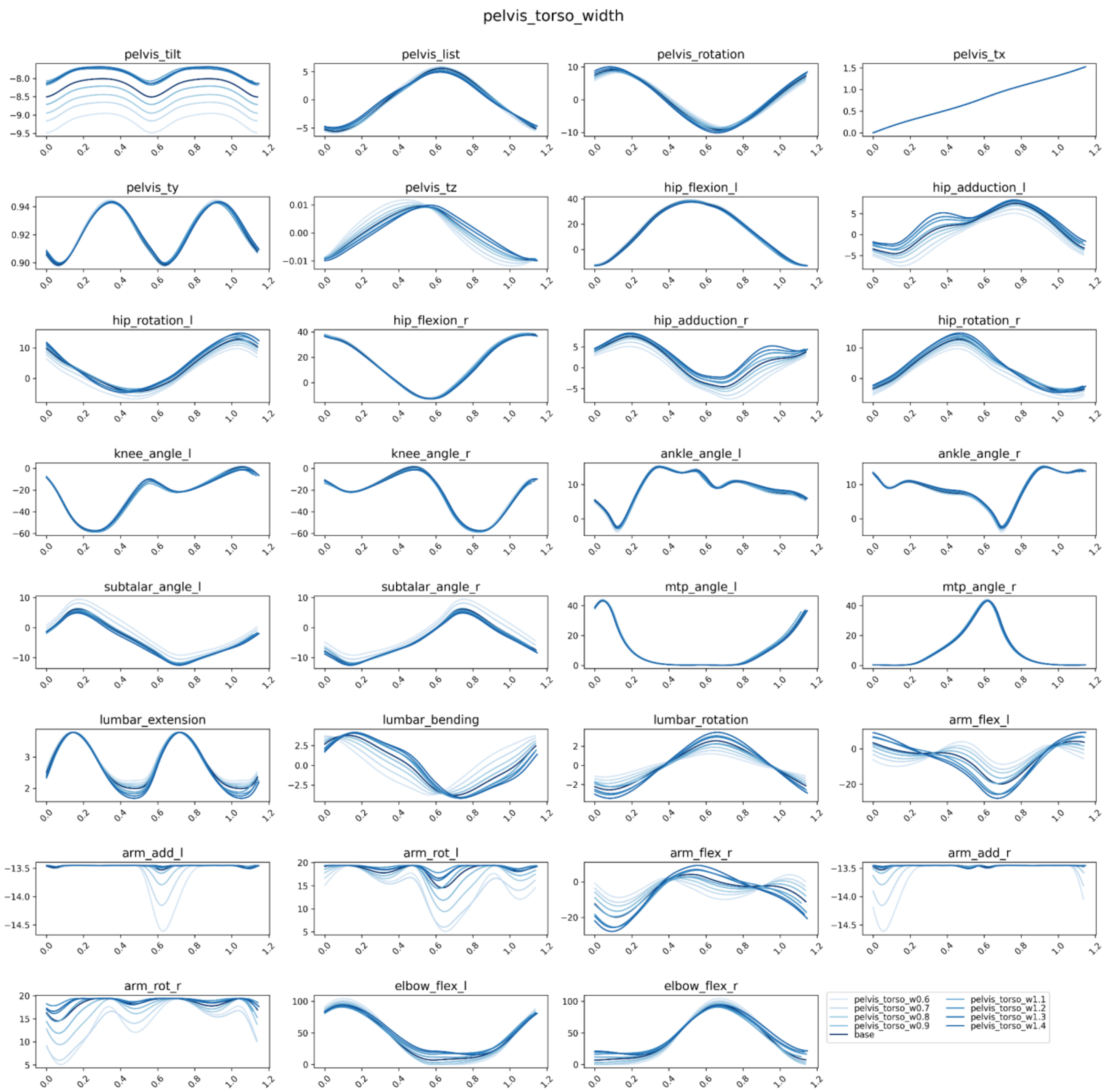
pelvis_torso_width
pelvis_tilt
pelvis_list
pelvis_rotation
pelvis_tx
pelvis_ty
pelvis_tz
hip_flexion_l
hip_adduction_l
hip_rotation_l
hip_flexion_r
hip_adduction_r
hip_rotation_r
knee_angle_l
knee_angle_r
ankle_angle_l
ankle_angle_r
subtalar_angle_l
subtalar_angle_r
mtp_angle_l
mtp_angle_r
lumbar_extension
lumbar_bending
lumbar_rotation
arm_flex_l
arm_add_l
arm_rot_l
arm_flex_r
arm_add_r
arm_rot_r
elbow_flex_l
elbow_flex_r
pelvis_torso_w0.6
pelvis_torso_w0.7
pelvis_torso_w0.8
pelvis_torso_w0.9
base
pelvis_torso_w1.1
pelvis_torso_w1.2
pelvis_torso_w1.3
pelvis_torso_w1.4

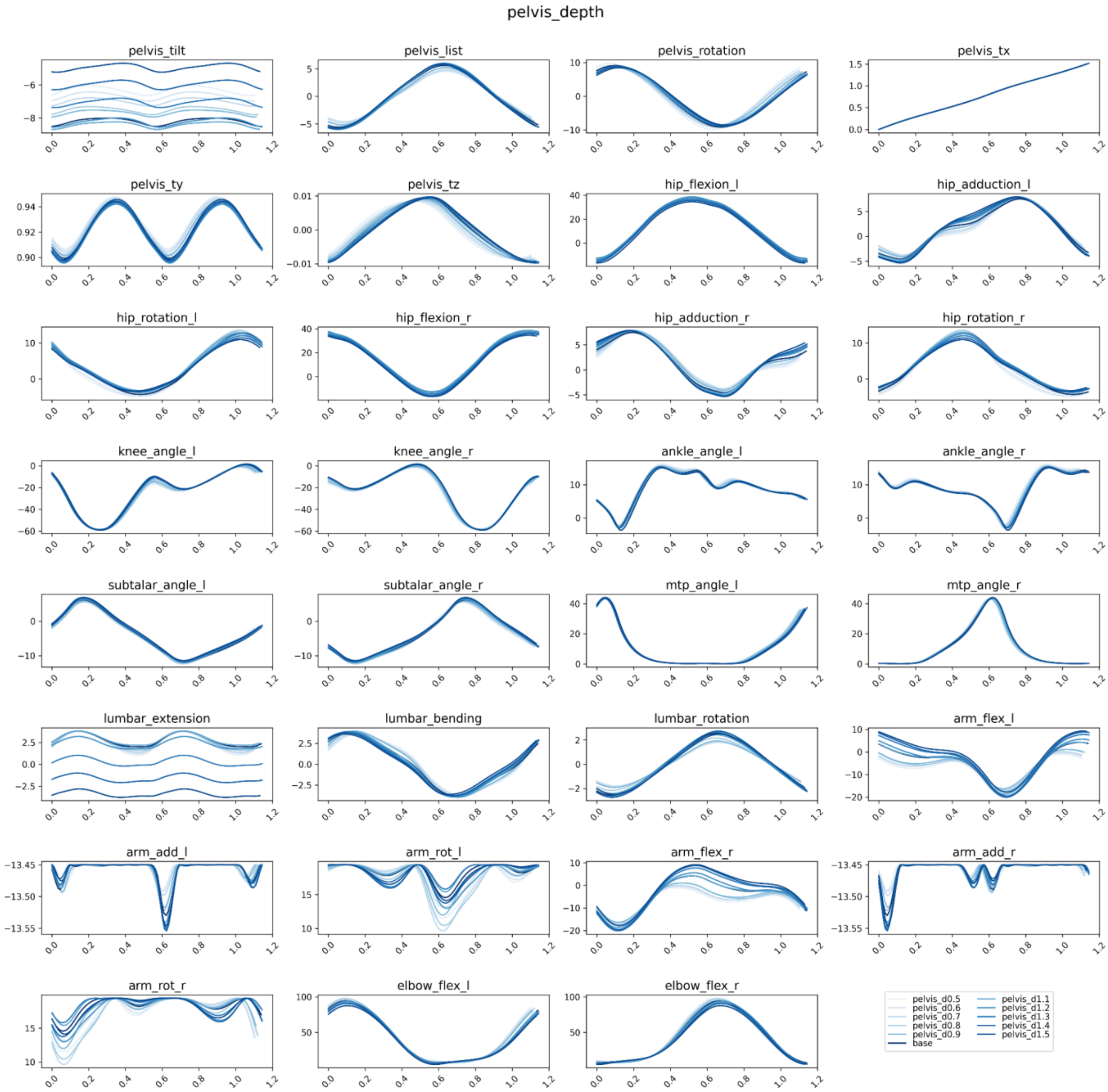
pelvis_depth
pelvis_tilt
pelvis_list
pelvis_rotation
pelvis_tx
pelvis_ty
pelvis_tz
hip_flexion_l
hip_adduction_l
hip_rotation_l
hip_flexion_r
hip_adduction_r
hip_rotation_r
knee_angle_l
knee_angle_r
ankle_angle_l
ankle_angle_r
subtalar_angle_l
subtalar_angle_r
mtp_angle_l
mtp_angle_r
lumbar_extension
lumbar_bending
lumbar_rotation
arm_flex_l
arm_add_l
arm_rot_l
arm_flex_r
arm_add_r
arm_rot_r
elbow_flex_l
elbow_flex_r
pelvis_d0.5
pelvis_d0.6
pelvis_d0.7
pelvis_d0.8
pelvis_d0.9
base
pelvis_d1.1
pelvis_d1.2
pelvis_d1.3
pelvis_d1.4
pelvis_d1.5

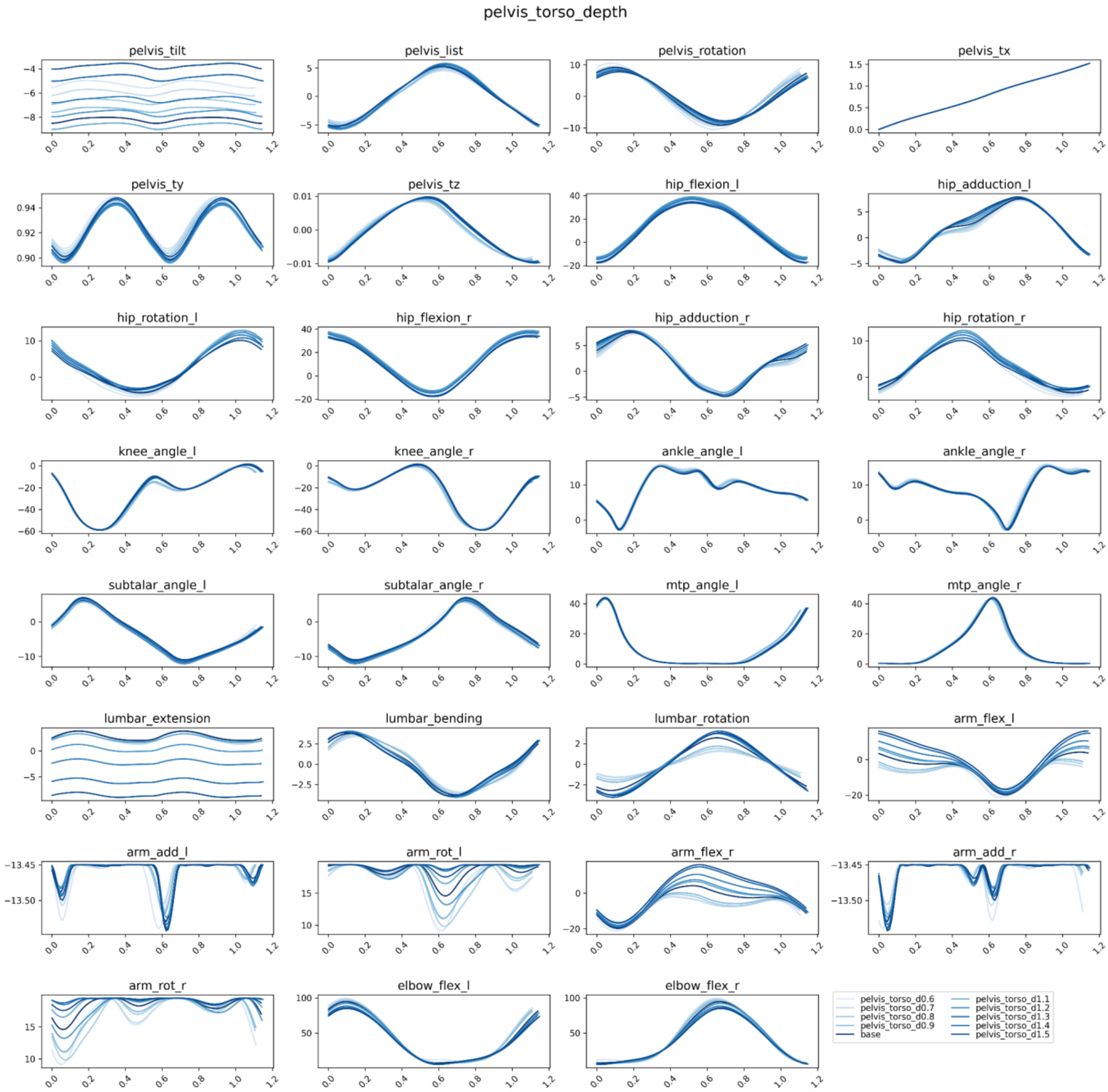

pelvis_torso_depth
pelvis_tilt
pelvis_list
pelvis_rotation
pelvis_tx
pelvis_ty
pelvis_tz
hip_flexion_l
hip_adduction_l
hip_rotation_l
hip_flexion_r
hip_adduction_r
hip_rotation_r
knee_angle_l
knee_angle_r
ankle_angle_l
ankle_angle_r
subtalar_angle_l
subtalar_angle_r
mtp_angle_l
mtp_angle_r
lumbar_extension
lumbar_bending
lumbar_rotation
arm_flex_l
arm_add_l
arm_rot_l
arm_flex_r
arm_add_r
arm_rot_r
elbow_flex_l
elbow_flex_r
pelvis_torso_d0.6
pelvis_torso_d0.7
pelvis_torso_d0.8
pelvis_torso_d0.9
base
pelvis_torso_d1.1
pelvis_torso_d1.2
pelvis_torso_d1.3
pelvis_torso_d1.4
pelvis_torso_d1.5

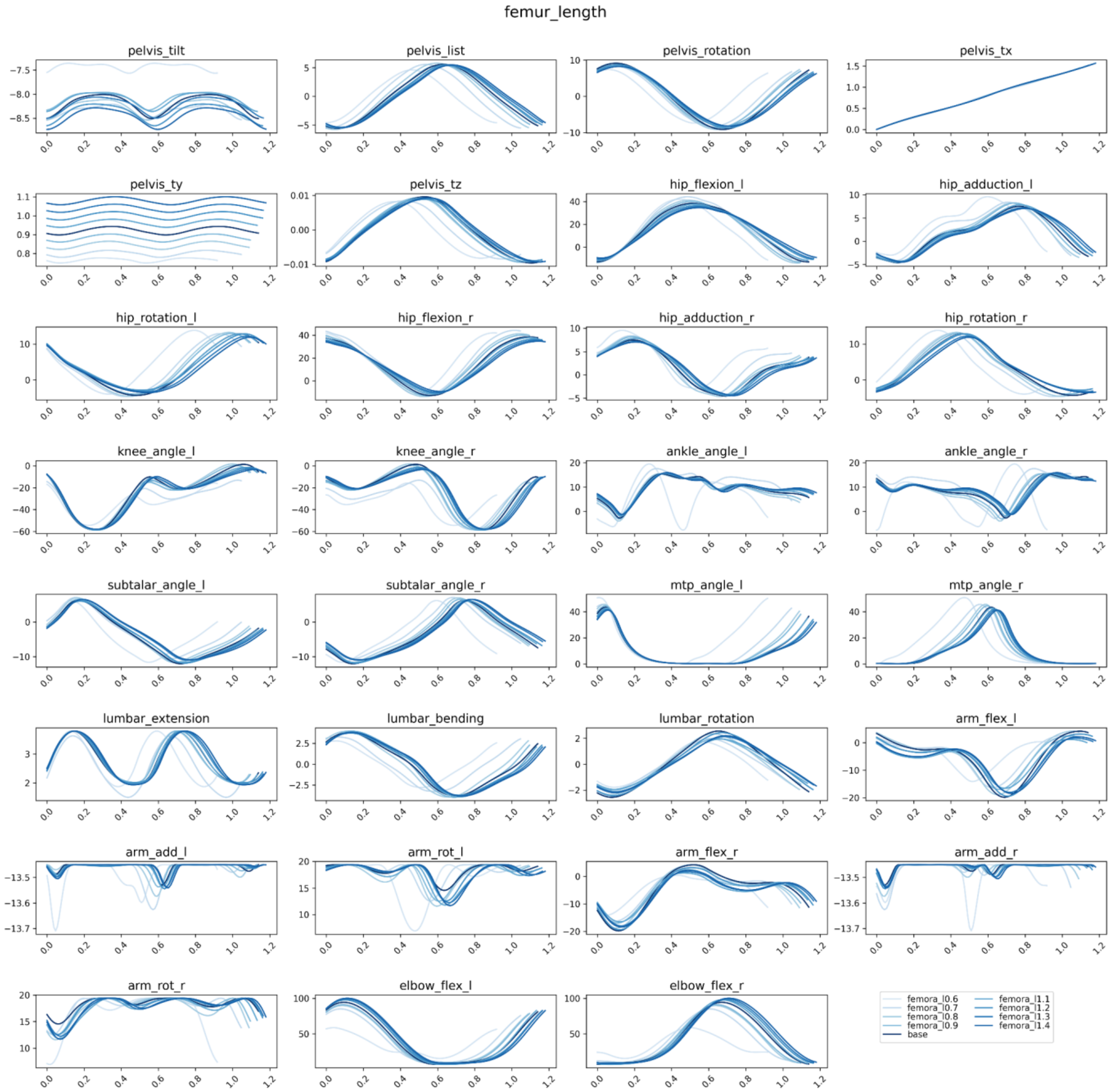
femur_length
pelvis_tilt
pelvis_list
pelvis_rotation
pelvis_tx
pelvis_ty
pelvis_tz
hip_flexion_l
hip_adduction_l
hip_rotation_l
hip_flexion_r
hip_adduction_r
hip_rotation_r
knee_angle_l
knee_angle_r
ankle_angle_l
ankle_angle_r
subtalar_angle_l
subtalar_angle_r
mtp_angle_l
mtp_angle_r
lumbar_extension
lumbar_bending
lumbar_rotation
arm_flex_l
arm_add_l
arm_rot_l
arm_flex_r
arm_add_r
arm_rot_r
elbow_flex_l
elbow_flex_r
femora_l0.6
femora_l0.7
femora_l0.8
femora_l0.9
base
femora_l1.1
femora_l1.2
femora_l1.3
femora_l1.4

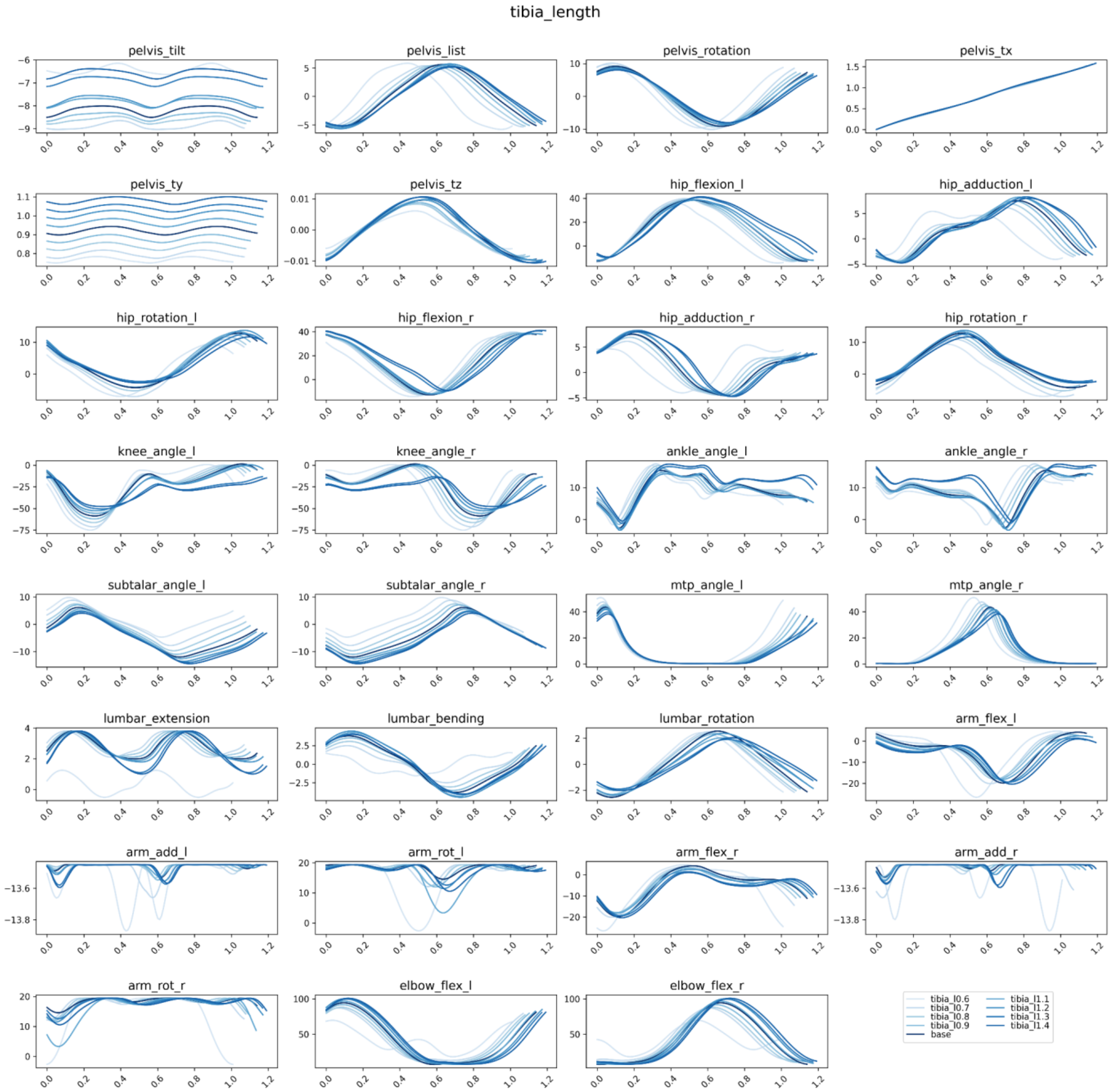
tibia_length
pelvis_tilt
pelvis_list
pelvis_rotation
pelvis_tx
pelvis_ty
pelvis_tz
hip_flexion_l
hip_adduction_l
hip_rotation_l
hip_flexion_r
hip_adduction_r
hip_rotation_r
knee_angle_l
knee_angle_r
ankle_angle_l
ankle_angle_r
subtalar_angle_l
subtalar_angle_r
mtp_angle_l
mtp_angle_r
lumbar_extension
lumbar_bending
lumbar_rotation
arm_flex_l
arm_add_l
arm_rot_l
arm_flex_r
arm_add_r
arm_rot_r
elbow_flex_l
elbow_flex_r
tibia_l0.6
tibia_l0.7
tibia_l0.8
tibia_l0.9
base
tibia_l1.1
tibia_l1.2
tibia_l1.3
tibia_l1.4

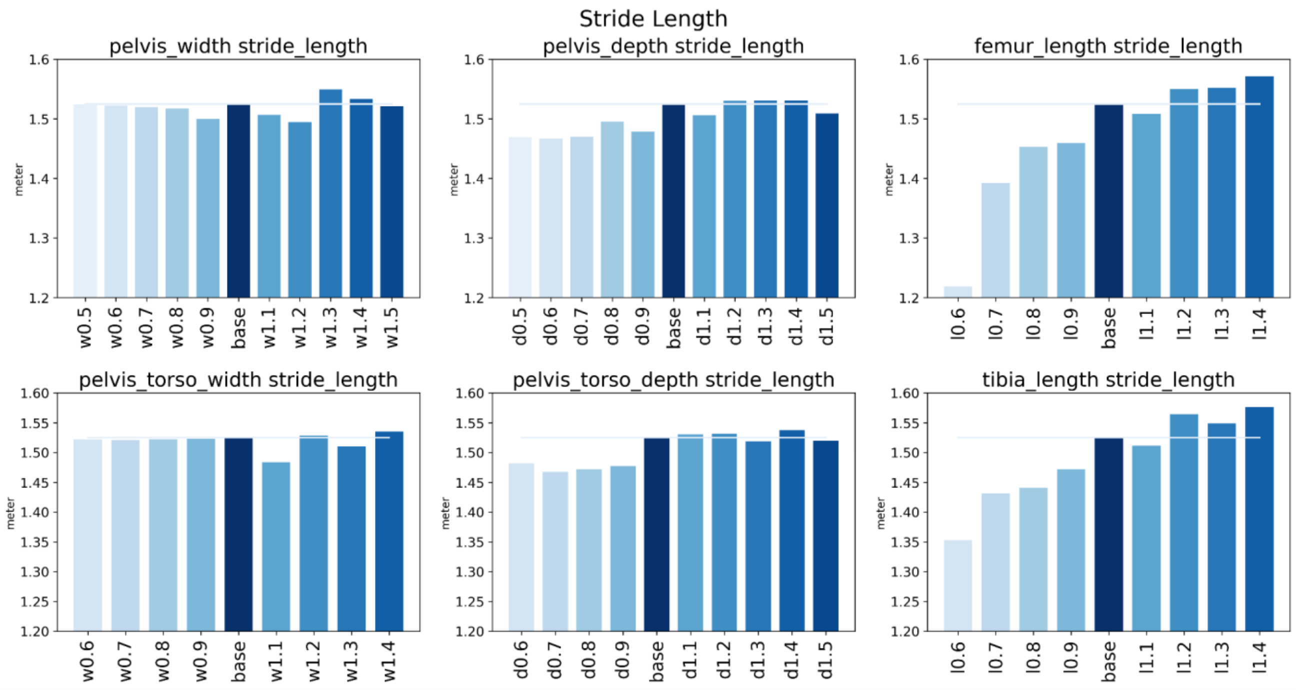


**Figure S2.** Stride length. Here, stride is two consecutive steps.

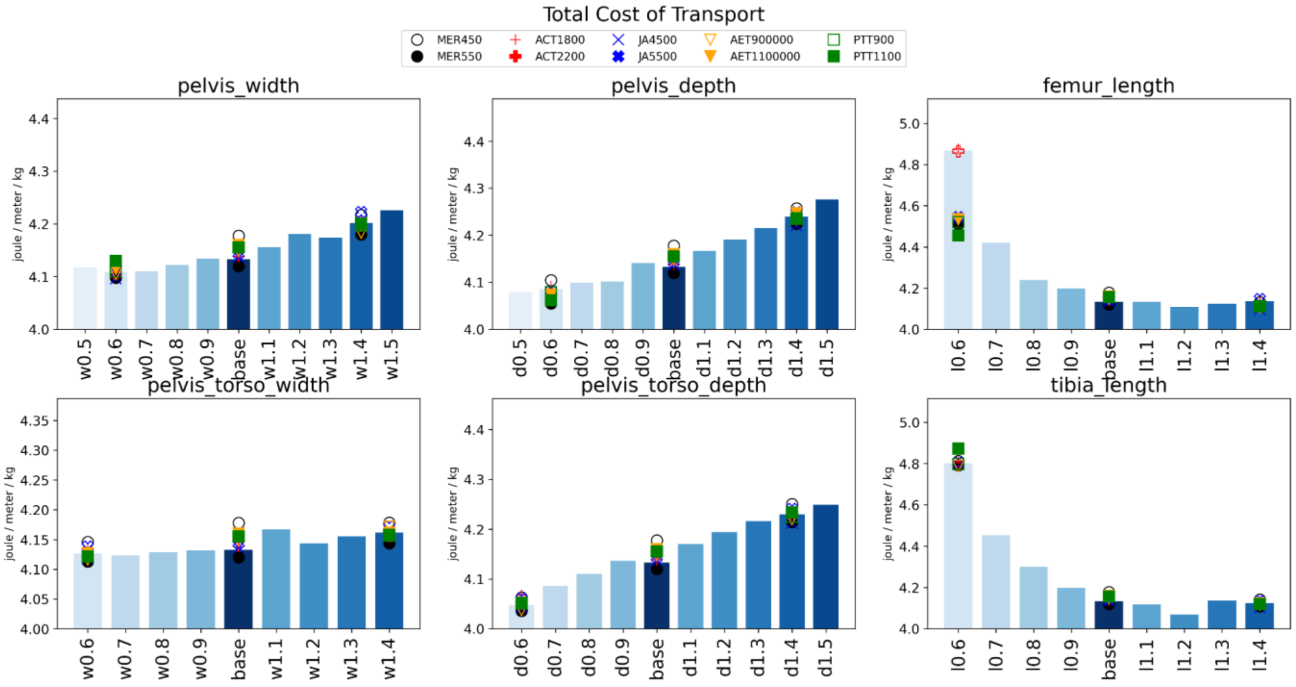


**Figure S3.** Variation of cost term weights:

- MER450 - Metabolic Energy Rate Term weight set at 450;
- MER550 - Metabolic Energy Rate Term weight set at 550;
- ACT1800 - Activation Term set at 1800;
- ACT2200 - Activation Term set at 2200;
- JA45000 - Joint Acceleration Term set at 45000;
- JA45000 - Joint Acceleration Term set at 55000;
- AET 900000 - Arm Excitation Term set at 900000;
- AET 1100000 - Arm Excitation Term set at 1100000;
- PTT900 - Passive Torque Term set at 900
- PTT1100 - Passive Torque Term set at 1100

Bars represent the results of the hot start trial with default values of weights:

- Metabolic Energy Rate Term: 450;
- Activation Term: 2000;
- Joint Acceleration Term: 50000;
- Arm Excitation Term: 1000000;
- Passive Torque Term: 1000.

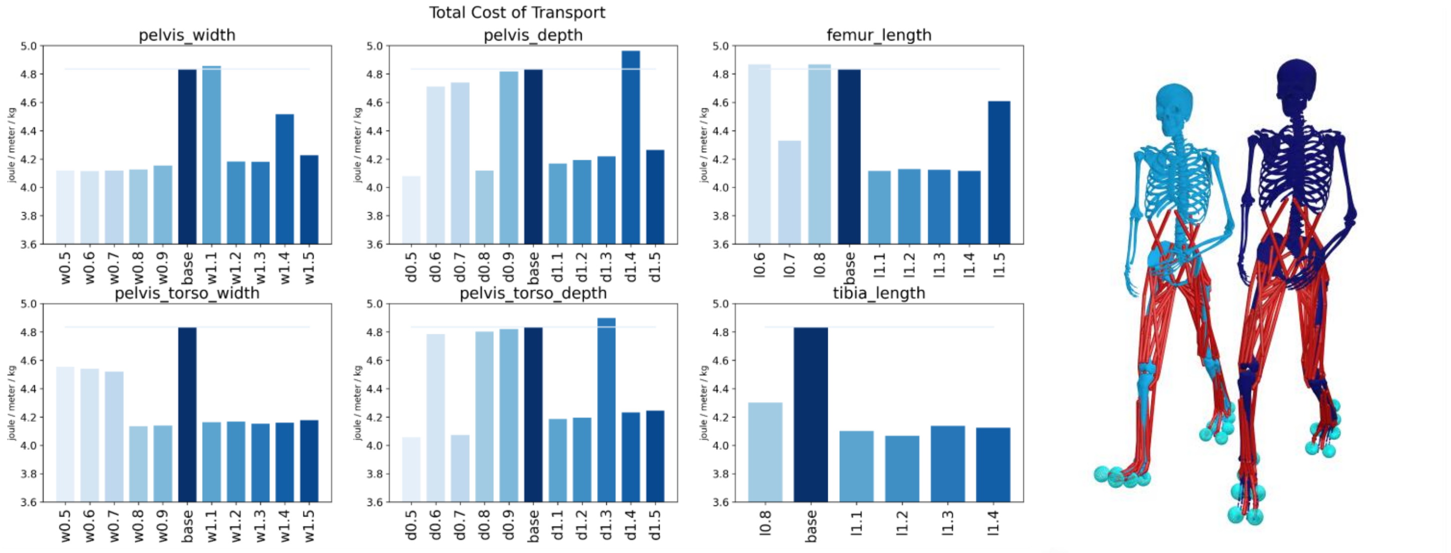


(a) (b)

**Figure S4.** (a) Cold start results for the total cost of transport. The initial guess was that all points of the time grid were set at 0; (b) Models with elevated COT produced a non-physiological walking gait on the toes. The model on the left has the tibia length scaled by a factor of 0.8. The model on the right is the base model.
